# Evolution of antibody immunity following Omicron BA.1 breakthrough infection

**DOI:** 10.1101/2022.09.21.508922

**Authors:** Chengzi I. Kaku, Tyler N. Starr, Panpan Zhou, Haley L. Dugan, Paul Khalifé, Ge Song, Elizabeth R. Champney, Daniel W. Mielcarz, James C. Geoghegan, Dennis R. Burton, Raiees Andrabi, Jesse D. Bloom, Laura M. Walker

**Author notes:** Corresponding author. (L.M.W.).

## Abstract

Understanding the evolution of antibody immunity following heterologous SAR-CoV-2 breakthrough infection will inform the development of next-generation vaccines. Here, we tracked SARS-CoV-2 receptor binding domain (RBD)-specific antibody responses up to six months following Omicron BA.1 breakthrough infection in mRNA-vaccinated individuals. Cross-reactive serum neutralizing antibody and memory B cell (MBC) responses declined by two- to four-fold through the study period. Breakthrough infection elicited minimal de novo Omicron-specific B cell responses but drove affinity maturation of pre-existing cross-reactive MBCs toward BA.1. Public clones dominated the neutralizing antibody response at both early and late time points, and their escape mutation profiles predicted newly emergent Omicron sublineages. The results demonstrate that heterologous SARS-CoV-2 variant exposure drives the evolution of B cell memory and suggest that convergent neutralizing antibody responses continue to shape viral evolution.

## Main text

The emergence and global spread of the SARS-CoV-2 Omicron BA.1 variant in late 2021 resulted in the largest surge in COVID-19 caseloads to date (*1*). While currently available COVID-19 vaccines induced high levels of protection against pre-Omicron variants, the extensive immune evasiveness of Omicron resulted in significantly reduced vaccine efficacy and durability following both primary and booster immunization (*2*–*5*). Moreover, antigenically drifted sub-lineages of Omicron (e.g. BA.2, BA.2.12.1, BA.4/5, BA.2.75, BA.2.75.2, and BA.4.6) continue to emerge and supplant prior sub-variants (*4*, *6*). The high prevalence of Omicron breakthrough infections led to the development and emergency use authorization of Omicron variant-based booster mRNA vaccines, despite limited immunogenicity and efficacy data in humans (*2*, *7*). Thus, there is an urgent need to understand if and how secondary exposure to antigenically divergent variants, such as Omicron, shape SARS-CoV-2-specific B cell memory.

We and others have previously reported that the acute antibody response following Omicron BA.1 breakthrough infection is dominated by re-activated memory B cells induced by mRNA vaccination (*8*–*11*). In support of these findings, preliminary data from clinical trials evaluating the immunogenicity of variant-based booster vaccines demonstrated that BA.1-containing mRNA vaccines induce a modest improvement in peak serum neutralizing responses compared with ancestral Wuhan-1 immunization (*12*). Although these studies provide evidence for “original antigenic sin” in the early B cell response following Omicron breakthrough infection, if and how this response evolves over time remains unclear. To address these questions, we longitudinally profiled SARS-CoV-2-specific serological and memory B responses in mRNA-vaccinated donors up to six months following BA.1 breakthrough infection.

We initially characterized the antibody response to SARS-CoV-2 in a cohort of seven mRNA-1273 vaccinated donors 14 to 27 days (median = 23 days) after BA.1 breakthrough infection (*8*). To study the evolution of this response, we obtained blood samples from six of the seven participants at a follow-up appointment four to six months (median = 153 days) post-infection (Fig. 1A, Table S1). Three of the six donors experienced infection after two-dose mRNA-1273 vaccination while the remaining three donors were infected after a third booster dose. None of the donors reported a second breakthrough infection between the two sample collection time points.

**Figure 1.**
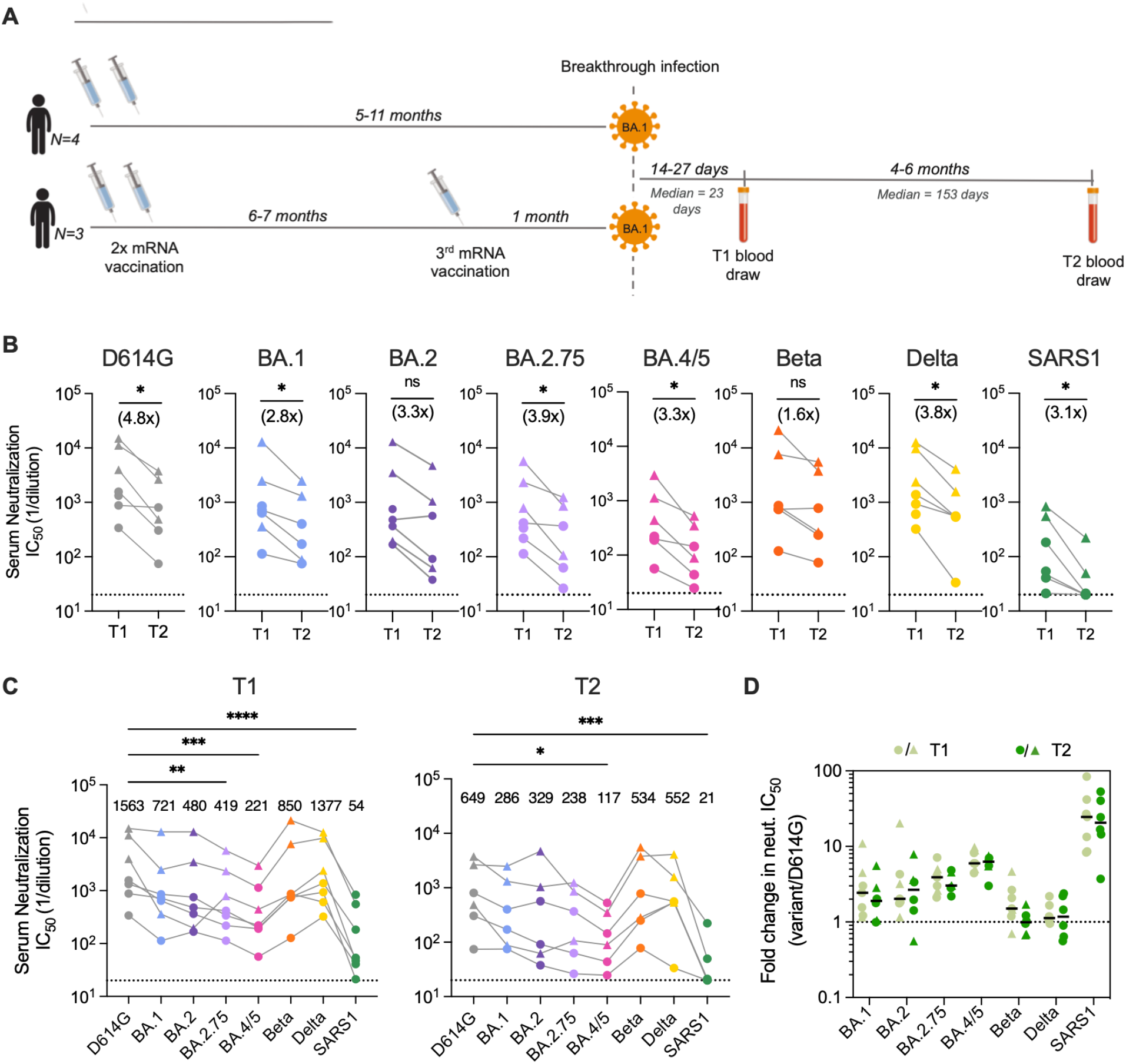
Serum neutralizing antibody responses induced following BA.1 breakthrough infection. **(A)** Timeline of vaccination, BA.1 breakthrough infection, and sample collections. **(B)** Paired analysis of serum neutralizing activity against SARS-CoV-2 D614G and BA.1, BA.2, BA.2.75, BA.4/5, Beta, and Delta variants, and SARS-CoV (SARS1) at 1-month (T1) and 5-6-month (T2) time points, as determined via a MLV-based pseudovirus neutralization assay. Connected data points represent paired samples for each donor, and the median fold change in serum titer between the two time points is shown in parentheses. Dotted lines represent the lower limit of detection of the assay. **(C)** Serum neutralizing titers against SARS-CoV-2 variants and SARS-CoV in samples collected at (left) 1-month and (right) 5-6-month post-breakthrough infection for each donor. Median titers are shown above the data points. Dotted lines represent the lower limit of detection of the assay. **(D)** Fold change in serum neutralizing titers for the indicated SARS-CoV-2 variants and SARS-CoV relative to SARS-CoV-2 D614G at early (T1) and late (T2) time points. Black bars represent median fold changes. Dotted line indicates no change in IC_50_. Breakthrough infection donors infected after two-dose mRNA vaccination (n = 4) are shown as circles and those infected after a third mRNA dose (n = 3) are shown as triangles. One two-dose vaccinated breakthrough donor was lost to follow-up at the second time point. Statistical comparisons were determined by (B) Wilcoxon matched-pairs signed rank test, (C) Friedman’s one-way ANOVA with Dunn’s multiple comparisons, or (D) mixed model ANOVA. \**P* < 0.05; \*\**P* < 0.01; \*\*\*\**P* < 0.0001; ns, not significant.

To evaluate serum neutralization breadth and potency, we tested the plasma samples for neutralizing activity against SARS-CoV-2 D614G, emergent variants (BA.1, BA.2, BA.4/5, BA.2.75, Beta, and Delta), and the more evolutionarily divergent sarbecovirus SARS-CoV, in a murine-leukemia virus (MLV)-based pseudovirus assay. Paired comparisons within each participant revealed that serum neutralizing titers against D614G declined by a median of 4.8-fold at 5- to 6-months post-infection relative to those observed within one-month post-infection (Fig. 1B). Correspondingly, we observed lower serum neutralizing titers against Omicron sub-variants (2.8 to 3.9-fold, respectively), Beta (1.6-fold), Delta (3.8-fold), and SARS-CoV (3.1-fold) at the 5- to 6-month time point relative to the early time point (Fig. 1B). Despite this waning of neutralizing antibody titers over time, all of the donor sera displayed detectable neutralizing activity against all of the SARS-CoV-2 variants tested at the 5-6 month time point (median titers ranging from 117 to 552) (Fig. 1C). Notably, titers remained within 3-fold of that observed for D614G for all variants except BA.4/5, which showed the greatest degree of escape from serum neutralizing antibodies (5.5-fold reduction from D614G), consistent with published serological studies (*4*, *5*). Furthermore, the fold reduction in serum neutralizing titer for SARS-CoV-2 VOCs relative to D614G remained similar at both time points, suggesting maintained serum neutralization breadth over time (Fig. 1D). We observed minimal cross-neutralizing activity against SARS-CoV (median titer = 21) in all donors, suggesting that serum neutralization breadth remained limited to SARS-CoV-2 variants (Fig. 1C). We conclude that serum neutralizing titers wane over the course 6-months following Omicron BA.1 breakthrough infection but nevertheless remain at detectable levels across a diverse range of SARS-CoV-2 variants through 6 months.

Next, we assessed the magnitude and cross-reactivity of the antigen-specific B cell response via flow cytometric enumeration of B cells stained with differentially labeled wildtype (Wuhan-1; WT) and BA.1 RBD tetramers (Fig. 2A, Fig. S1A). At the 5-6-month time point, total RBD-reactive B cells (WT and/or BA.1-reactive) and WT/BA.1 cross-reactive B cells comprised a median of 0.44% (ranging 0.12-2.53%) and 0.37% (ranging 0.12-2.53%) of class-switched (IgG^+^ or IgA^+^) B cells, respectively (Fig. 2B, 2C, Fig. S1A). Thus, 86% (ranging 69-100%) of all RBD+ class-switched B cells at 5-6 months post-infection displayed BA.1/WT cross-reactivity, compared with 75% at 1-month post-infection (ranging 65-81%) (Fig. 2D, Fig. S2). Correspondingly, WT-specific B cells decreased from 25% of all RBD+ class-switched B cells at 1 month to 11% at 5-6 months (Fig. 2D, Fig. S2). Consistent with the waning of serum neutralizing titers over time, we also observed a modest but significant decline (1.1 to 3.7-fold) in the frequencies of WT/BA.1 cross-reactive B cells at 5-6 months relative to the 1-month time point (Fig. 2C). At the late time point, we also detected the emergence of a BA.1-specific B cell population (average = 3% of class-switched B cells) in 3 of the 6 individuals, although the magnitude of this response varied widely among individuals (ranging from 1-18%) (Fig. 2D, Fig. S2). In summary, Omicron BA.1 breakthrough infection induces a WT/BA.1 cross-reactive B cell response at early time points post-infection and this response only modestly declines over the course of 6 months.

**Figure 2.**
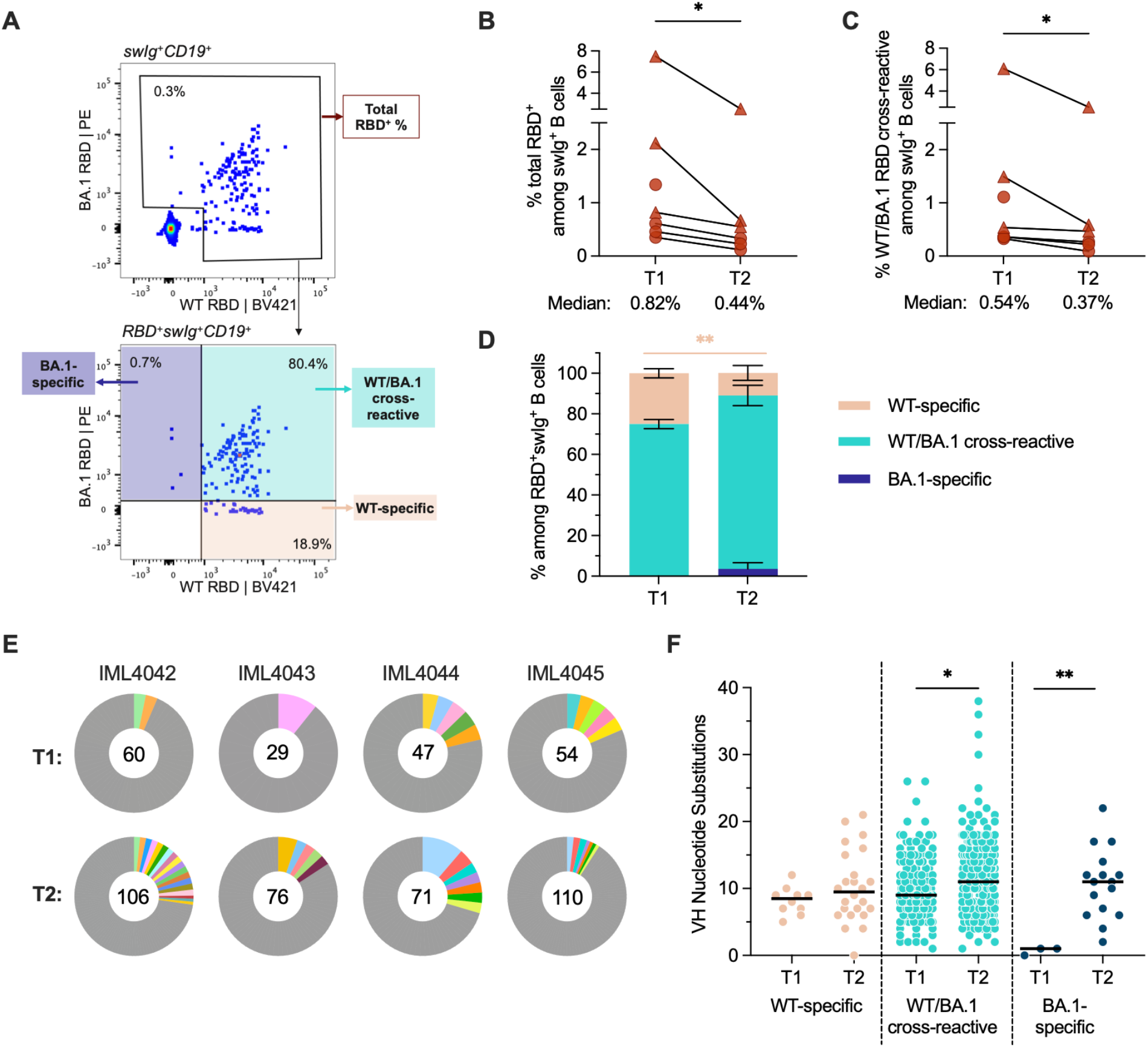
SARS-CoV-2 RBD-specific memory B cell responses following BA.1 breakthrough infection. **(A)** Representative fluorescence-activated cell sorting gating strategy used to enumerate frequencies of (top) total (WT+BA.1) RBD-reactive B cells among class-switched (IgG^+^ or IgA^+^) CD19^+^ B cells and (bottom) WT-specific, BA.1-specific, and WT/BA.1 cross-reactive B cells among total RBD-reactive, class-switched (IgG^+^ or IgA^+^) CD19^+^ B cells. **(B-C)** Frequencies of (B) total RBD-reactive or (C) WT/BA.1 RBD cross-reactive B cells among class-switched CD19^+^ B cells at 1-month (T1) and 5-6-month (T2) time points. Connected data points represent paired samples for each donor. Donors infected after two-dose mRNA vaccination (n = 4) are shown as circles and those infected after a third mRNA dose (n = 3) are shown as triangles. One two-dose vaccinated breakthrough donor was censored at the second time point. **(D)** Mean proportions of RBD-reactive, class-switched B cells that display WT-specific, BA.1-specific or WT/BA.1-cross-reactive binding at each time point. Error bars indicate standard error of mean. **(E)** Clonal lineage analysis of RBD-directed antibodies isolated from four donors at the early (T1) and late (T2) time points. Clonally expanded lineages (defined as antibodies with the same heavy and light chain germlines, same CDR3 lengths, and > 80% CDRH3 sequence identity) are represented as colored slices. Each colored slice represents a clonal lineage with the size of the slice proportional to the lineage size. Unique clones are combined into a single gray segment. The number of antibodies is shown in the center of each pie. Three of the donors (IML4042, IML4043, and IML4044) experienced BA.1 breakthrough infection following two-dose mRNA vaccination and the remaining donor (IML4045) was infected after a booster immunization. **(F)** Levels of somatic hypermutation, as determined by the number of nucleotide substitutions in the variable heavy (VH) region, at the early and late time points among WT-specific, WT/BA.1 cross-reactive, and BA.1-specific antibodies. Medians are shown by black bars. Statistical significance was determined by (B and C) Wilcoxon matched-pairs signed rank test or (D and F) Mann-Whitney U test. swIg^+^, class-switched immunoglobulin. PE, phycoerythrin; \**P* < 0.05; \*\**P* < 0.01.

To compare the molecular characteristics of antibodies isolated at early and late time points following BA.1 breakthrough infection, we single-cell sorted 71 to 110 class-switched RBD-reactive B cells from four of the five previously studied donors (donors IML4042, IML4043, IML4044, IML4045) at 139 to 170 days after breakthrough infection and expressed a total of 363 natively paired antibodies as full-length IgGs (Fig. S1B) (*8*). Similar to the antibodies characterized from the acute time point, the newly isolated antibodies primarily recognized both WT and BA.1 RBD antigens (73-97%), exhibited a high degree of clonal diversity, and displayed preferential usage of certain VH germline genes (*IGHV1-46, 1-69, 3-13, 3-53, 3-66, 3-9*, and *4-31* germline genes at both time points) (Fig. 2E, Fig. S3 and S4). The level of SHM in the cross-reactive antibodies increased from a median of 9 VH nucleotide substitutions at 1-month to 11 VH nucleotide substitutions by 5-6 months, potentially suggesting affinity maturation in secondary germinal centers (Fig. 2F). Consistent with their higher levels of SHM, the antibodies isolated at 5-6 months displayed 1.7-fold improved binding to BA.1 (median K_D_ = 1.3 nM) and 2-fold reduced binding affinity to the WT RBD (median K_D_ = 1.0 nM) relative to early antibodies, suggesting maturation towards Omicron BA.1 at the expense of WT affinity (Fig. 3A and 3B). These changes in binding recognition resulted in the late antibodies showing more balanced affinity profiles compared to the early antibodies (Fig 3B). For example, the majority of antibodies (73%) isolated at the late time point exhibited WT and BA.1 RBD affinities within two-fold of each other compared to only 24% of early antibodies (Fig. 3C).

**Figure 3.**
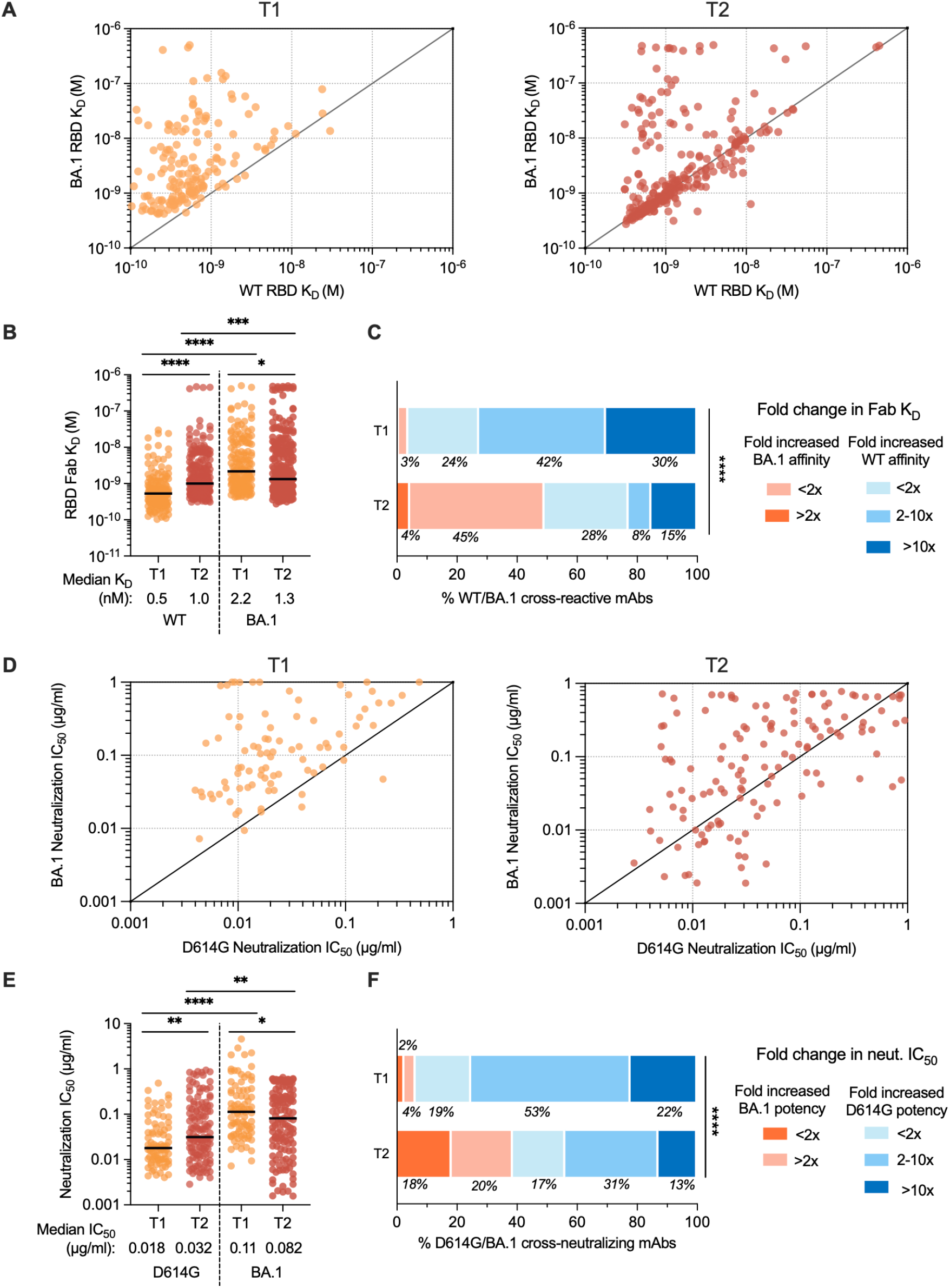
Binding and neutralizing properties of RBD-directed antibodies induced by BA.1 breakthrough infection. **(A-B)** Fab binding affinities of WT/BA.1 cross-reactive antibodies for recombinant WT and BA.1 RBD antigens, as measured by BLI, are plotted as bivariates for antibodies derived from (left) 1-month and (right) 5-6-month time points in (A) and summarized as a column dot plot in (B). Median affinities are indicated by black bars and shown below data points. **(C)** Proportions of WT/BA.1 cross-reactive antibodies at each time point that show increased affinity for the BA.1 RBD relative to WT (red shades) or increased affinity for WT RBD (blue shades). Values represent the percentage of antibodies belonging to each of the indicated categories. **(D-E)** Neutralizing activities of cross-binding antibodies against SARS-CoV-2 D614G and BA.1, as determined by an MLV-based pseudovirus neutralization assay. IC_50_ values are plotted in (D) as bivariates for antibodies isolated from (left) 1-month and (right) 5-6-month tie points and summarized as column dot plots in (E). Median IC_50_ values are indicated by black bars and shown below data points. **(F)** Proportions of WT/BA.1 cross-neutralizing antibodies at each time point that show increased neutralizing potency against BA.1 (red shades) or D614G (blue shades). Values represent the percentage of antibodies belonging to each of the indicated categories. Statistical comparisons were determined by (B and E) multiple Mann-Whitney U tests without adjustment for multiplicity across time points and Wilcoxon matched-pairs rank tests within each time point or (C and F) Mann-Whitney U test. IC_50_, 50% inhibitory concentration; K_D_, equilibrium dissociation constant; \**P* < 0.05; \*\**P* < 0.01; \*\*\*\**P* < 0.0001.

To determine whether the improvement in binding affinity for BA.1 translated into enhanced neutralization potency, we assessed the antibodies for neutralizing activity against WT and BA.1 using a pseudovirus assay. Fifty-one percent and 42% of WT/BA.1 cross-binding antibodies isolated from the 1-month and 5-6-month time point, respectively, cross-neutralized D614G and BA.1 with IC_50_ < 2 μg/ml. Overall, the neutralizing antibodies displayed approximately 2-fold lower potency against D614G at the late time point relative to the acute time point, consistent with the observed reduction in WT RBD affinity over time (Fig. 3D and 3E). As expected, the improvement in BA.1 binding affinities over time translated into an overall improvement in neutralization potency (Fig. 3E). Approximately 16% of antibodies isolated at 5-6 months displayed neutralization IC_50_s <0.01 ug/ml compared to only 2% of antibodies isolated at the earlier time point (Fig. S5). As a result, forty-one percent of the neutralizing antibodies isolated at 6 months exhibited more potent activity against BA.1 relative to D614G, compared to only 7% of the acute neutralizing antibodies (Fig 3F). In summary, cross-reactive antibody responses induced following BA.1 breakthrough infection evolve toward increased BA.1 affinity and neutralization potency for at least 6 months post-infection.

Although the vast majority of antibodies isolated at the 5-6-month time point displayed WT/BA.1 cross-reactive binding, we identified a limited number of BA.1-specific antibodies in all four donors, comprising 1% to 15% of total RBD-specific antibodies (median = 4%) (Fig. S3). In contrast, we only detected BA.1-specific antibodies in a single donor at the acute time point (Fig. S3). Furthermore, unlike the BA.1-specific antibodies isolated at the early time point, which lacked somatic mutations, the BA.1-specific antibodies identified at 5-6 months displayed SHM levels similar to those of cross-reactive antibodies (median = 11 VH nucleotide substitutions) (Fig. 2F). Forty percent of BA.1-specific antibodies isolated at the late time point neutralized BA.1, with IC_50_s ranging from 0.002 to 0.089 μg/ml, and none of the antibodies displayed detectable neutralizing activity against D614G (Fig. S6). Thus, BA.1 breakthrough infection induces a limited and delayed de novo Omicron-specific B cell response that undergoes affinity maturation over time.

To further explore the breadth of both WT/BA.1 cross-reactive and BA.1-specific neutralizing antibodies, we evaluated their binding reactivities with a panel of recombinant RBDs encoding mutations present in SARS-CoV-2 variants BA.2, BA.4/5, Beta, and Delta, and the more antigenically divergent SARS-CoV. D614G/BA.1 cross-neutralizing antibodies displayed 2.4-fold reduced affinity for the WT RBD and 3.4-fold improved affinity for the BA.1 RBD relative to early neutralizing antibodies, consistent with the pattern observed for all WT/BA.1 cross-binding antibodies (Fig. 4A, Fig. 3A-C). Furthermore, the WT/BA.1 cross-reactive antibodies isolated at 6 months broadly recognized other SARS-CoV-2 variants, except for BA.4/5, which was associated with a ≥5-fold loss in affinity for 57% (68/120) of the WT/BA.1 neutralizing antibodies (Fig. 4A, Fig. S7). Importantly, the 5-6-month antibodies displayed higher affinity binding to all Omicron sub-variants and Beta relative to the early antibodies, suggesting that the increased affinity to BA.1 also improved breadth of reactivity against other variants (Fig. 4A). In support of this finding, a significantly higher proportion (40%) of neutralizing antibodies isolated at 6 months displayed high affinity (K_D_ < 10nM) binding to all five variants tested compared with early antibodies (22%) (Fig. 4B). Furthermore, antibodies isolated at the late time point displayed smaller differences in binding affinity against BA.1, BA.2, BA.4/5 and the early Beta and Delta variants relative to early antibodies (Fig. 4C). In contrast to the WT/BA.1 cross-reactive antibodies, the BA.1-specific neutralizing antibodies displayed limited breadth, with only 50% of these antibodies maintaining binding to BA.2 and none of the antibodies showing reactivity with WT, BA.4/5, Beta, or Delta (Fig. S6). We conclude that BA.1 breakthrough infection results in an overall broadening of the anti-SARS-CoV-2 neutralizing antibody repertoire.

**Figure 4.**
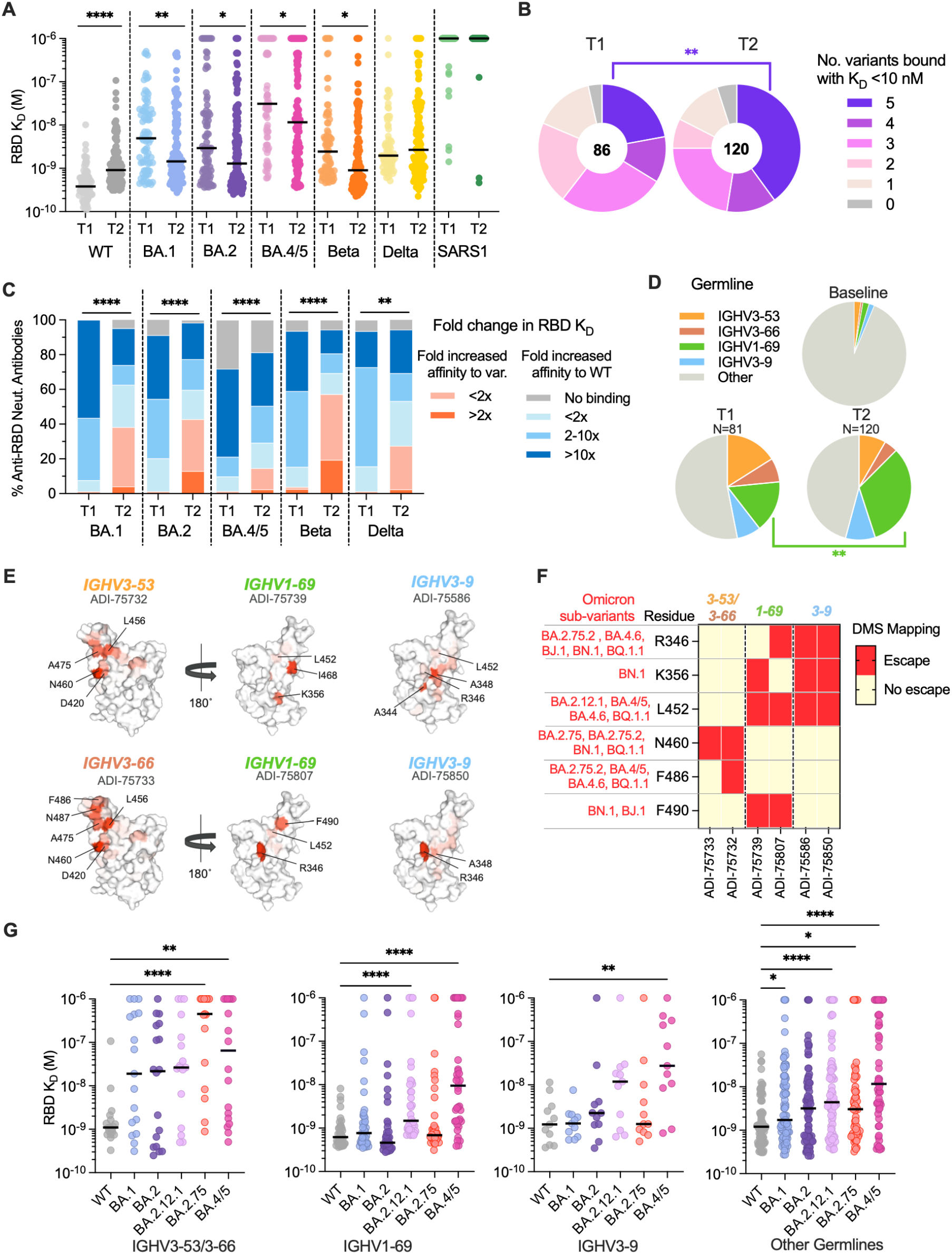
Breadth of D614G/BA.1 cross-neutralizing antibodies at early and late time points following BA.1 breakthrough infection. **(A)** Fab binding affinities of D614G/BA.1 cross-neutralizing antibodies isolated at 1-month (T1) and 5-6-month (T2) time points for recombinant SARS-CoV-2 variant RBDs and the SARS-CoV RBD, as determined by BLI. Black bars represent medians. **(B)** Pie charts showing the proportions of antibodies derived from (left) early and (right) late time points that bound the indicated number of SARS-CoV-2 variant RBDs with Fab K_D_s < 10 nM. The total number of antibodies is shown in the center of each pie. **(C)** Proportions of D614G/BA.1 cross-neutralizing antibodies with the indicated fold changes in Fab binding affinities for recombinant SARS-CoV-2 variant RBDs relative to the WT RBD. **(D)** Pie charts showing frequencies of the indicated convergent germline genes among D614G/BA.1 cross-neutralizing antibodies isolated at early (T1) and late (T2) timelines. Germline gene frequencies observed in baseline human antibody repertoires (upper right) are shown for comparison (*25*). **(E)** Structural projections of binding escape mutations determined for the indicated convergent antibodies using deep mutational scanning analysis of yeast-displayed SARS-CoV-2 BA.1 RBD mutant libraries. The RBD surface is colored by a gradient ranging from no escape (white) to strong escape (red) at each site. See Fig. S10 for additional details. **(F)** Heatmap summarizing convergent antibody escape mutations present in the indicated SARS-CoV-2 Omicron sub-lineages. **(G)** Fab binding affinities of convergent antibodies utilizing the indicated germline genes for SARS-CoV-2 WT and Omicron sub-variant RBD antigens, as measured by BLI. Black bars indicate median affinities. Statistical comparisons were determined by (A and C) Kruskal-Wallis test with Holms corrected multiple pairwise comparisons, (B and D) Fisher’s exact test, or (G) Kruskal-Wallis test with subsequent Dunn’s multiple comparisons with WT. K_D_, equilibrium dissociation constant; \**P* < 0.05; \*\**P* < 0.01; \*\*\**P* < 0.001; \*\*\*\**P* < 0.0001.

Among neutralizing antibodies isolated at both time points, we observed significant over-representation of four IGHV germline genes (*IGHV1-69, IGHV3-53/3-66*, and *IGHV3-9*) (*8*) (Fig S8A). At the 5-6-month time point, over half (54%) of the neutralizing antibodies were encoded by one of these four germlines, with one-third of these antibodies utilizing *IGHV1-69* (Fig. 4D, Fig. S8). We previously found that BA.1-neutralizing *IGHV1-69* antibodies isolated from the early time point preferentially paired with the light chain germline IGLV1-40 and targeted an antigenic site overlapping that of the class 3 antibody COV2-2130 and non-overlapping with the ACE2 binding site (*8*). Similarly, 69% of *IGHV1-69* antibodies isolated at 5-6 months paired with the *IGLV1-40* germline and the majority (80%) failed to compete with ACE2 for binding (Fig. S9A and C). Likewise, >90% of *IGHV3-9* antibodies identified from both time points recognized a non-ACE2-competitive binding site, although unlike *IGHV1-69* antibodies, *IGHV3-9* antibodies recognize an epitope overlapping S309 and REGN10987 as well as COV2-2130, suggesting a distinct mode of binding from IGHV1-69 antibodies (Fig. S9C) (*8*). Lastly, *IGHV3-53/66* antibodies isolated from both time points were characterized by short HCDR3s (median = 11 to 12 nucleotide substitutions) compared with baseline HCDR3 lengths (median = 15 substitutions) and displayed competitive binding with the ACE2 receptor (Fig. S9B and C). Thus, convergent antibody classes dominated the neutralizing antibody response at both early and late time points following BA.1 breakthrough infection, suggesting little to no change in B cell immunodominance hierarchy over time.

Given the dominance of these public clonotypes in BA.1 breakthrough infection donors, we sought to determine their escape mutations in the BA.1 background. We randomly selected one to two antibodies belonging to each convergent germline and performed deep mutational scanning (DMS) analysis using a library encoding all possible amino acid substitutions from BA.1 (Fig. S10A) (*13*). Antibodies encoded by *IGHV3-53* (ADI-75733) and *IGHV3-66* (ADI-75732) displayed similar escape profiles, consistent with their shared sequence features and competitive binding profiles (Fig. 4E and Fig. S10C) (*8*). RBD positions N460 and F486, which are mutated in emergent variants (N460K in B.2.75, BA.2.75.2, BN.1, and BQ.1; F486S in BA.2.75.2; and F486V in BA.4/5, BA.4.6, and BQ.1.1), were associated with binding escape from *IGHV3-53/66* antibodies (Fig. 4F and Fig. 10C). *IGHV1-69* and *IGHV3-9* antibodies both showed reduced binding to RBDs incorporating mutations at positions 344-349, 356, 452-453, 468, and 490. Notably, residues R346, K356, L452, and F490 are mutated across evolutionarily diverse Omicron sub-lineages, including BA.4.6 (R346T, L452R), BA.4/5 (L452R), BA.2.12.1 (L452Q), BJ.1 (R346T, F490V), BN.1 (R346T, K356T, F490S), and BQ.1.1 (R346T, L452R) (Fig. 4F and Fig. S10C). Consistent with these escape profiles, *IGHV1-69* and *IGHV3-9* class antibodies displayed reduced binding to BA.2.12.1 and BA.4/5 relative to early Omicron variants, likely due to the unique L452Q/R mutations present in these variants compared with BA.1 and BA.2 (Fig. 4G). Consistent with DMS-based predictions, both BA.2.75 and BA.4/5 RBDs displayed increased binding resistance to *IGHV3-53/66* antibodies (Fig. 4F and 4G). Thus, convergent D614G/BA.1 cross-neutralizing antibodies recognize epitopes commonly mutated in recently emerging Omicron sub-variants, providing a molecular explanation for the high degree of antigenic convergence observed in recent Omicron sub-variant evolution and their increased level of immune evasion relative to BA.1.

In summary, BA.1 breakthrough infection in mRNA-vaccinated individuals induces broadly neutralizing serological and MBC responses that persist for at least six months after infection, supporting real-world studies showing that BA.1 breakthrough infection provides protection against symptomatic BA.1, BA.2, and BA.5 infection for at least 5-6 months (*14*–*16*).

Furthermore, although the acute B cell response following breakthrough infection is primarily mediated by recall of cross-reactive vaccine-induced MBCs, these MBC clones accumulate somatic mutations and evolve increased breadth and potency for at least 6 months following infection. Although this enhanced neutralization breadth and potency was not reflected in the serum antibody response, it is possible that a second heterologous exposure may broaden the serological repertoire by activating these affinity matured MBCs, akin to the improved serum neutralization breadth observed following mRNA booster vaccination (*17*, *18*). Nevertheless, our data indicate that infection or vaccination with antigenically divergent SARS-CoV-2 variants may provide long-term benefits by broadening pre-existing anti-SARS-CoV-2 B cell memory.

Finally, we found that convergent classes of neutralizing antibodies dominated the BA.1 breakthrough response at both early and late time points, reminiscent of the antibody response elicited following primarily infection or vaccination with early ancestral SARS-CoV-2 strains (*19*–*21*). The sustained prevalence of public clones that target residues frequently mutated in emerging Omicron subvariants suggests that this response is the driving force behind the continued antigenic drift of Omicron. Thus, in contrast to current approaches to the design of universal vaccines for certain highly antigenically variable viruses, such as HIV and influenza, which aim to focus the neutralizing response on a limited number of relatively conserved epitopes, the development of “variant-proof” COVID-19 vaccines may require a different strategy: engineering of spike-based immunogens that induce a diversity of neutralizing antibodies targeting numerous co-dominant epitopes, with the goal of limiting convergent immune pressure and therefore constraining viral evolution (*22*–*24*).

## Acknowledgements

We thank T. Boland for assistance with sequence analysis. We acknowledge E. Krauland, J. Nett, M. Vasquez, and C.G. Rappazzo for helpful comments on the manuscript. We also thank the Flow Cytometry and Genomics Shared Facilities at Fred Hutchinson Cancer Center. All IgGs were sequenced by Adimab’s Molecular Core and produced by the High-Throughput Expression group.

## Author contributions

C.I.K. and L.M.W. conceived and designed the study. D.M. supervised and performed clinical sample collection and processing. C.I.K. designed and performed B cell analyses. C.I.K. and P.K. performed single B cell sorting. C.I.K., P.Z., P.K., and H.L.D. performed pseudovirus neutralization assays. T.N.S. designed and performed antibody deep mutational scanning analyses. C.I.K. and E.R.C. performed biolayer interferometry assays. C.I.K., H.L.D., and G.S. conducted antibody sequence analyses. C.I.K., T.N.S., H.L.D., J.C.G., D.R.B., R.A., J.D.B., and L.M.W. analyzed the data. C.I.K. and L.M.W. wrote the manuscript, and all authors reviewed and edited the paper.

## Data and materials availability

Omicron BA.1 yeast-display deep mutational scanning libraries are available from Addgene (accession # 1000000187). Complete computational pipeline with intermediate and final data files is available from GitHub: https://github.com/jbloomlab/SARS-CoV-2-RBD_Omicron_MAP_Adimab. All other data needed to evaluate the conclusions in the paper are present in the paper or the Supplementary Materials. IgGs are available from L.M.W. under a material transfer agreement from Invivyd Inc.

## Supplementary Materials

Materials and Methods

Figures S1 – S10

Table S1

References 25 – 30

## Supplementary Materials

### Materials and Methods

#### Human subjects and blood sample collection

Seven BA.1 breakthrough infected participants were recruited to participate in this study with informed consent under the healthy donor protocol D10083, Immune Monitoring Core (DartLab) Laboratory at Dartmouth-Hitchcock Hospital, as previously described (*8*). Briefly, participants experienced breakthrough infection after two- or three-dose mRNA vaccination (BNT162b2 and/or mRNA-1273). Venous blood was collected at two time points, an early visit at 14 to 27 days (T1) and a late visit 139 to 170 days (T2) after their first SARS-CoV-2 test. Participants had no documented history of SARS-CoV-2 infection prior to vaccination or between the two blood draw time points. Clinical and demographic characteristics of breakthrough infection donors are shown in Table S1. Plasma and peripheral blood mononuclear cell (PBMC) samples were isolated using a Ficoll 1077 (Sigma) gradient, as previously described (*8*).

#### Plasmid Design and Construction

Plasmids expressing spike proteins of SARS-CoV-2 variants and SARS-CoV were ordered as gene block fragments (IDT) and cloned into a mammalian expression vector for MLV-based pseudovirus production as previously described (*26*). All SARS-CoV-2 variant spikes and the SARS-CoV spike were C-terminally truncated by 19-amino acids or 28-amino acids, respectively, to increase infectious titers. The SARS-CoV S sequence was retrieved from ENA (AAP13441). SARS-CoV-2 variants contain the following mutations from the Wuhan-Hu-1 sequence (Genbank: NC_045512.2):

- D614G: D614G
- Beta: D80A, D215G, Δ242-244, K417N, E484K, N501Y, D614G, A701V
- Delta: T19R, G142D, Δ156-157, R158G, L452R, T478K, D614G, P681R, D950N
- BA.1: A67V, Δ69-70, T95I, G142D/Δ143-145, Δ211/L212I, ins214EPE, G339D, S371L, S373P, S375F, K417N, N440K, G446S, S477N, T478K, E484A, Q493R, G496S, Q498R, N501Y, Y505H, T547K, D614G, H655Y, N679K, P681H, N764K, D796Y, N856K, Q954H, N969K, L981F
- BA.2: T19I, L24S, Δ25-27, G142D, V213G, G339D, S371F, S373P, S375F, T376A, D405N, R408S, K417N, N440K, S477N, T478K, E484A, Q493R, Q498R, N501Y, Y505H, D614G, H655Y, N679K, P681H, N764K, D796Y, Q954H, N969K
- BA.4/5: T19I, L24S, Δ25-27, Δ69-70, G142D, V213G, G339D, S371F, S373P, S375F, T376A, D405N, R408S, K417N, N440K, L452R, S477N, T478K, E484A, F486V, Q498R, N501Y, Y505H, D614G, H655Y, N679K, P681H, N764K, D796Y, Q954H, N969K
- BA.2.75: T19I, L24S, Δ25-27, G142D, K147E, W152R, F157L, I210V, V213G, G339H, G257S, S371F, S373P, S375F, T376A, D405N, R408S, K417N, N440K, G446S, N460K, S477N, T478K, E484A, Q498R, N501Y, Y505H, D614G, H655Y, N679K, P681H, N764K, D796Y, Q954H, N969K

#### SARS-CoV-2 pseudovirus generation

Single-cycle infectious MLVs pseudotyped with spike proteins of SARS-CoV-2 variants and SARS-CoV were generated as previously described (*26*). Briefly, HEK293T cells were seeded at a density of 0.5 million cells/ml in 6-well tissue culture plates and the next day, transfected using Lipofectamine 2000 (ThermoFisher Scientific) with the following plasmids: 1) 0.5 μg per well of pCDNA3.3 encoding SARS-CoV-2 spike with a 19-amino acid truncation at the C-terminus, 2) 2 μg per well of MLV-based luciferase reporter gene plasmid (Vector Builder), and 3) 2 μg per well of of MLV gag/pol (Vector Builder). MLV particles were harvested 48 h post-transfection, aliquoted, and stored at −80 °C for neutralization assays.

#### Pseudovirus neutralization assay

MLV pseudovirus neutralization assays for serum and monoclonal antibodies were performed as previously described (*8*). Briefly, 56 °C heat-inactivated sera or antibodies were serially diluted in 50 μl MEM/EBSS media supplemented with 10% fetal bovine serum (FBS) and incubated with 50 μl of MLV viral stock for 1 h at 37 °C. Following incubation, antibody-virus mixtures were added to previously seeded HeLa-hACE2 reporter cells (BPS Bioscience Cat #79958). Infection was allowed to occur for 48 h at 37 °C. Infection was measured by lysing cells with Luciferase Cell Culture Lysis reagent (Promega) and detecting luciferase activity using the Luciferase Assay System (Promega) following manufacturer’s protocols. Infectivity was as quantified by relative luminescence units (RLUs) and the percentage neutralization was calculated as 100*(1–[RLU_sample_– RLU_background_]/[RLU_isotype control mAb_–RLU_background_]). Neutralization IC_50_ was interpolated from curves fitted using four-parameter non-linear regression in GraphPad Prism (version 9.3.1).

#### FACS analysis of SARS-CoV-2 S-specific B cell responses

Antigen-specific B cells were detected using recombinant biotinylated antigens tetramerized with fluorophore-conjugated streptavidin (SA), as previously described (*8*). Briefly, Avitag biotinylated WT RBD (Acro Biosystems, Cat #SPD-C82E8) and Avitag biotinylated BA.1 RBD (Acro Biosystems, Cat # SPD-C82E4) were mixed in 4:1 molar ratios with SA-BV421 (BioLegend) and SA-phycoerythrin (PE; Invitrogen), respectively, and allowed to incubate for 20 min on ice. Unbound SA sites were subsequently quenched using 5 μl of 2 μM Pierce biotin (ThermoFisher Scientific). Approximately 10 million PBMCs were stained with tetramerized RBDs (25 nM each); anti-human antibodies anti-CD19 (PE-Cy7; Biolegend), anti-CD3 (PerCP-Cy5.5; Biolegend), anti-CD8 (PerCP-Cy5.5; Biolegend), anti-CD14 (PerCP-Cy5.5; Invitrogen), and anti-CD16 (PerCP-Cy5.5; Biolegend); and 50 μl Brilliant Stain Buffer (BD BioSciences) diluted in FACS buffer (2% BSA/1 mM EDTA in 1X PBS). 200 μl of staining reagents were added to each PBMC sample and incubated for 15 min on ice. After one wash with FACS buffer, cells were stained in a mixture of propidium iodide and anti-human antibodies anti-IgG (BV605; BD Biosciences), anti-IgA (FITC; Abcam), anti-CD27 (BV510; BD Biosciences), and anti-CD71 (APC-Cy7; Biolegend). Following 15 min of incubation on ice, cells were washed two times with FACS buffer and analyzed using a BD FACS Aria II (BD BioSciences).

For sorting of RBD-specific, class-switched B cells, PBMCs that react with either WT and/or BA.1 RBD tetramers among CD19^+^CD3^−^CD8^−^CD14^−^CD16^−^PI^−^ and IgG^+^ or IgA^+^ cells were single-cell index sorted into 96-well polystyrene microplates (Corning) containing 20 μl lysis buffer per well [5 μl of 5X first strand SSIV cDNA buffer (Invitrogen), 1.25 μl dithiothreitol (Invitrogen), 0.625 μl of NP-40 (Thermo Scientific), 0.25 μl RNaseOUT (Invitrogen), and 12.8 μl dH2O]. Plates briefly centrifuged and then frozen at −80 °C before PCR amplification.

#### Amplification and cloning of antibody variable genes

Antibody variable gene fragments (VH, Vk, Vλ) were amplified by RT-PCR as described previously (*27*). Briefly, cDNA was synthesized using randomized hexamers and SuperScript IV enzyme (ThermoFisher Scientific). cDNA was subsequently amplified by two rounds of nested PCRs, with the second cycle of nested PCR adding 40 base pairs of flanking DNA homologous to restriction enzyme-digested *S. cerevisiae* expression vectors to enable homologous recombination during transformation. PCR-amplified variable gene DNA was mixed with expression vectors and chemically transformed into competent yeast cells via the lithium acetate method (*28*). Yeast were plated on selective amino acid drop-out agar plates and individual yeast colonies were picked for sequencing and recombinant antibody expression.

#### Expression and purification of IgG and Fab molecules

Antibodies were expressed as human IgG1 via *S. cerevisiae* cultures, as described previously (*27*). Briefly, yeast cells were grown in culture for 6 days for antibody production, before collecting IgG-containing supernatant by centrifugation. IgGs were subsequently purified by protein A-affinity chromatography and eluted using 200 mM acetic acid/50 mM NaCl (pH 3.5). The pH was then neutralized using 1/8^th^ volume of 2 M Hepes (pH 8.0). Fab fragments were cleaved from full-length IgG by incubating with papain for 2 h at 30 °C before terminating the reaction using iodoacetamide. Fab fragments were purified from the mixture of digested antibody Fab ad Fc fragments using a two-step chromatography system: 1) Protein A agarose was used to remove Fc fragments and undigested IgG, and 2) Fabs in the flow-through were further purified using CaptureSelect™ IgG-CH1 affinity resin (ThermoFisher Scientific) and eluted from the column using 200 mM acetic acid/50 mM NaCl (pH 3.5). Fab solutions were pH-neutralized using 1/8th volume 2 M Hepes (pH 8.0).

#### Binding affinity measurements by biolayer interferometry

Antibody binding kinetics were measured by biolayer interferometry (BLI) using a FortéBio Octet HTX instrument (Sartorius). All steps were performed at 25 °C and at an orbital shaking speed of 1000 rpm, and all reagents were formulated in PBSF buffer (PBS with 0.1% w/v BSA). To measure monovalent binding affinities against SARS-CoV-2 RBD variants and SARS-CoV S, recombinant RBDs of SARS-CoV-2 WT (Acro Biosystems, Cat #SPD-C52H3), Beta (Acro Biosystems, Cat #SPD-C52Hp), Delta (Acro Biosystems, Cat #SPD-C52Hh), BA.1 (Acro Biosystems, Cat #SPD-C522f), BA.2 (Acro Biosystems, Cat#SPD-C522g), BA.4/5 (Acro Biosystems, Cat#SPD-C522r), and SARS-CoV (Sino Biological, Cat #40150-V08B2) were biotinylated using EZ-Link™ Sulfo-NHS-LC-Biotin (Thermo Scientific) following manufacturer’s recommendations to achieve an average of 4 biotins per RBD molecule. Biotinylated antigens were diluted (100 nM) in PBSF and loaded onto streptavidin biosensors (Sartorius) to a sensor response of 1.0-1.2 nm and then allowed to equilibrate in PBSF for a minimum of 30 min. After a 60 s baseline step in PBSF, antigen-loaded sensors were exposed (180 s) to 100 nM Fab and then dipped (420 s) into PBSF to measure any dissociation of the antigen from the biosensor surface. Fab binding data with detectable binding responses (>0.1 nm) were aligned, inter-step corrected (to the association step) and fit to a 1:1 binding model using the FortéBio Data Analysis Software (version 11.1).

#### ACE2 competition by biolayer interferometry

Antibody binding competition with recombinant human ACE2 receptor (Sino Biological, Cat# 10108-H08H) was determined by BLI using a ForteBio Octet HTX (Sartorius). All binding steps were performed at 25 °C and at an orbital shaking speed of 1000 rpm. All reagents were formulated in PBSF (1X PBS with 0.1% w/v BSA). IgGs (100 nM) were captured onto anti-human IgG capture (AHC) biosensors (Molecular Devices) to a sensor response of 1.0 nm-1.4 nm, and then soaked (20 min) in an irrelevant IgG1 solution (0.5 mg/ml) to block remaining Fc binding sites. Next, sensors were equilibrated for 30 min in PBSF and then briefly exposed (90 s) to 300 nM of ACE2 to assess any potential cross interactions between sensor-loaded IgG and ACE2. Sensors were allowed to baseline (60 s) in PBSF before exposing (180 s) to 100 nM SARS-CoV-2 RBD (Acro Biosystems, Cat # SPD-C52H3). Last, RBD-bound sensors were exposed (180 s) to 300 nM ACE2 to assess competition, where antibodies that resulted in increased sensor responses after ACE2 exposure represented non-ACE2-competitive binding profiles while those resulting in unchanged responses represented ACE2-competitive profiles.

#### Deep mutational scanning analysis of antibody binding escape

Yeast-display deep mutational scanning experiments identifying mutations that escape binding by each monoclonal antibody were conducted with duplicate site-saturation mutagenesis Omicron BA.1 RBD libraries (*13*). Yeast libraries were grown in SD-CAA media (6.7 g/L Yeast Nitrogen Base, 5.0 g/L Casamino acids, 2.13 g/L MES, and 2% w/v dextrose), and backdiluted to 0.67 OD600 in SG-CAA+0.1%D (SD-CAA with 2% galactose and 0.1% dextrose in place of the 2% dextrose) to induce RBD expression, which proceeded for 16-18 hours at room temperature with mild agitation. 5 OD of cells were washed in PBS-BSA (0.2 mg/L) and labeled for one hour at room temperature in 1 mL with a concentration of antibody determined as the EC90 from pilot isogenic binding assays. In parallel, 0.5 OD of yeast expressing the Omicron BA.1 wildtype RBD were incubated in 100 μL of antibody at the matched EC90 concentration or 0.1x the concentration for FACS gate-setting. Cells were washed, incubated with 1:100 FITC-conjugated chicken anti-Myc antibody (Immunology Consultants CMYC-45F) to label RBD expression and 1:200 PE-conjugated goat anti-human-IgG (Jackson ImmunoResearch 109-115-098) to label bound antibody. Labeled cells were washed and resuspended in PBS for FACS.

Antibody-escape cells in each library were selected via FACS on a BD FACSAria II. FACS selection gates were drawn to capture approximately 50% of yeast expressing the wildtype BA.1 RBD labeled at 10x reduced antibody labeling concentration (see gates in Fig. S10A). For each sample, ~4 million RBD^+^ cells were processed on the sorter with collection of cells in the antibody-escape bin. Sorted cells were grown overnight in SD-CAA + pen-strep, plasmid purified (Zymo D2005), PCR amplified, and barcode sequenced on an Illumina NextSeq. In parallel, plasmid samples were purified from 30 OD of pre-sorted library cultures and sequenced to establish pre-selection barcode frequencies.

Demultiplexed Illumina barcode reads were matched to library barcodes in barcode-mutant lookup tables using dms_variants (version 0.8.9), yielding a table of counts of each barcode in each pre- and post-sort population which is available at https://github.com/jbloomlab/SARS-CoV-2-RBD_Omicron_MAP_Adimab/blob/main/results/counts/variant_counts.csv.gz. The escape fraction of each barcoded variant was computed from sequencing counts in the pre-sort and antibody-escape populations via the formula:

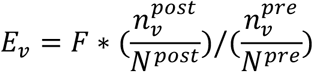

where *F* is the total fraction of the library that escapes antibody binding, *n*_v_ is the counts of variant *v* in the pre- or post-sort samples with a pseudocount addition of 0.5, and *N* is the total sequencing count across all variants pre- and post-sort. These escape fractions represent the estimated fraction of cells expressing a particular variant that fall in the escape bin. Per-barcode escape scores are available at https://github.com/jbloomlab/SARS-CoV-2-RBD_Omicron_MAP_Adimab/blob/main/results/escape_scores/scores.csv.

We applied computational filters to remove mutants with low sequencing counts or highly deleterious mutations that had ACE2 binding scores < –2 or expression scores of < –1, and we removed mutations to the conserved RBD cysteine residues. Per-mutant escape fractions were computed as the average across barcodes within replicates, with the correlations between replicate library selections shown in Fig. S10B. Final escape fraction measurements averaged across replicates are available at https://github.com/jbloomlab/SARS-CoV-2-RBD_Omicron_MAP_Adimab/blob/main/results/supp_data/Adimabs_raw_data.csv.

## Supplementary Figures

**Fig. S1.**
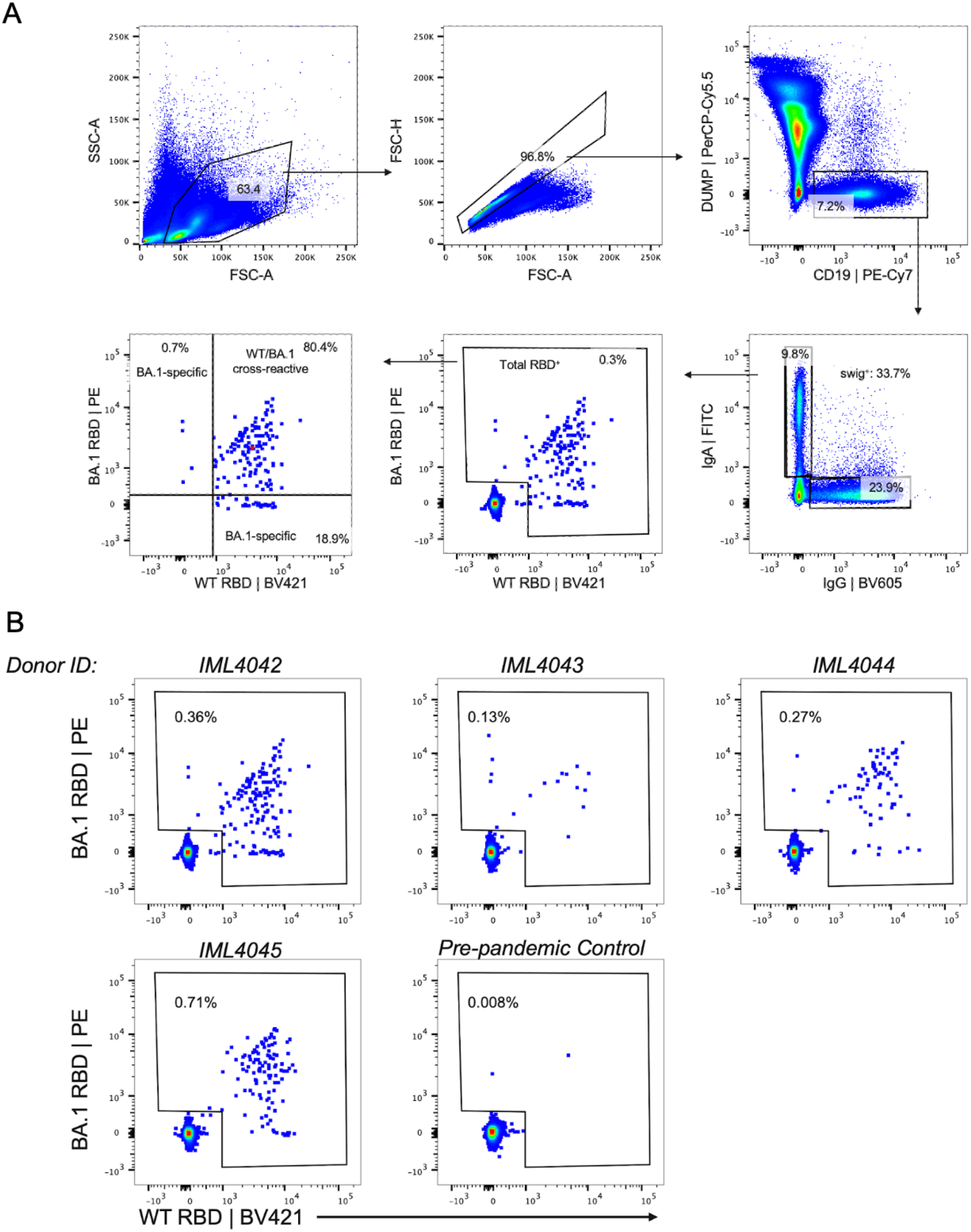
SARS-CoV-2 antigen-specific B cell staining and sorting. **(A)** Representative FACS gating strategy to determine frequencies of WT and/or BA.1 RBD-reactive B cells among class-switched (IgG^+^ or IgA^+^) B cells. The frequency of events in each gate relative to the parent gate are shown as percentages in each plot. **(B)** FACS gates used for single-cell sorting of WT and/or Omicron BA.1 RBD-specific memory B cells in 4 individuals 5-6 months following BA.1 breakthrough infection, Donors IML4042, IML4043, and IML4044 experienced breakthrough infection following two-dose mRNA vaccination, and IML4045 was infected after a third mRNA dose. A healthy pre-pandemic donor sample is shown as a control. FSC-A, forward scatter area; FSC-H, forward scatter height; swIg^+^, class-switched immunoglobulin; SSC-A, side scatter area.

**Fig. S2.**
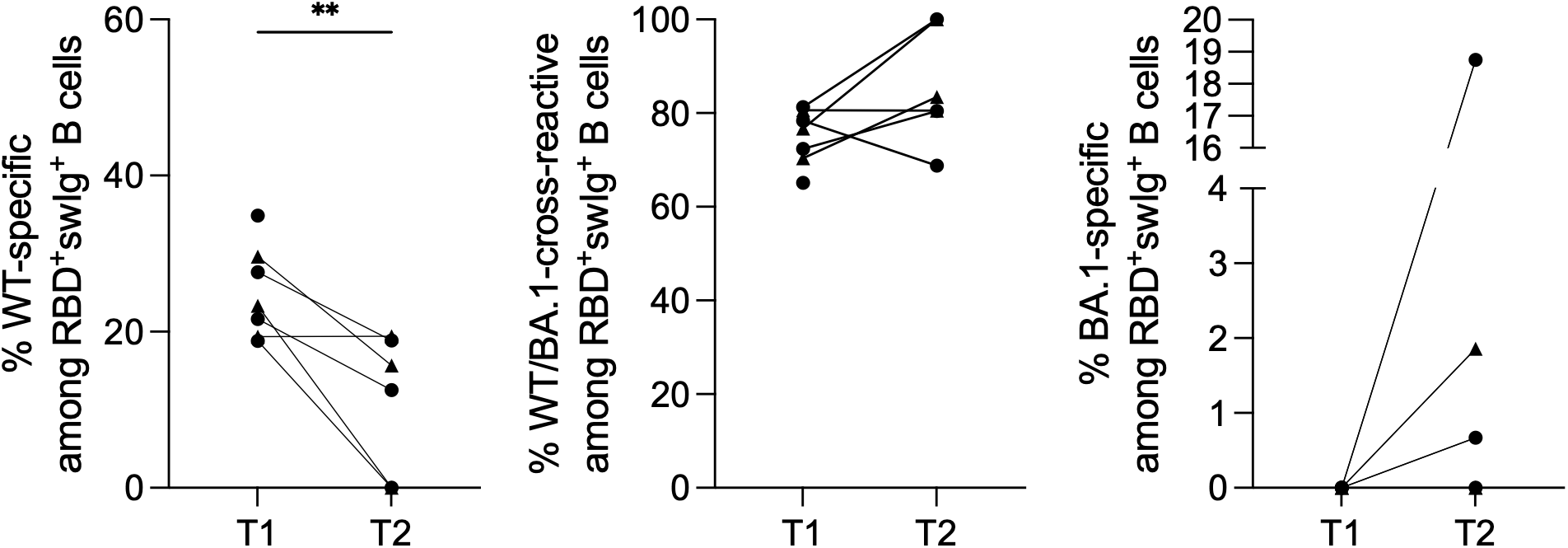
Cross-reactivity of RBD-directed B cells at early and late time points following BA.1 breakthrough infection. Proportion of RBD-directed class-switched B cells that are (left) WT-specific, (middle) WT/BA.1 cross-reactive, and (right) BA.1-specific at 1-month (T1) and 5-6-month (T2) time points, as determined by flow cytometry. Donors infected after two-dose mRNA vaccination (n = 4) are shown as circles and those infected after a third mRNA booster dose (n = 3) are shown as triangles. One two-dose vaccinated breakthrough donor was censored at the second time point. Statistical comparisons were determined by Mann-Whitney U tests. \*\**P* < 0.01.

**Fig. S3.**
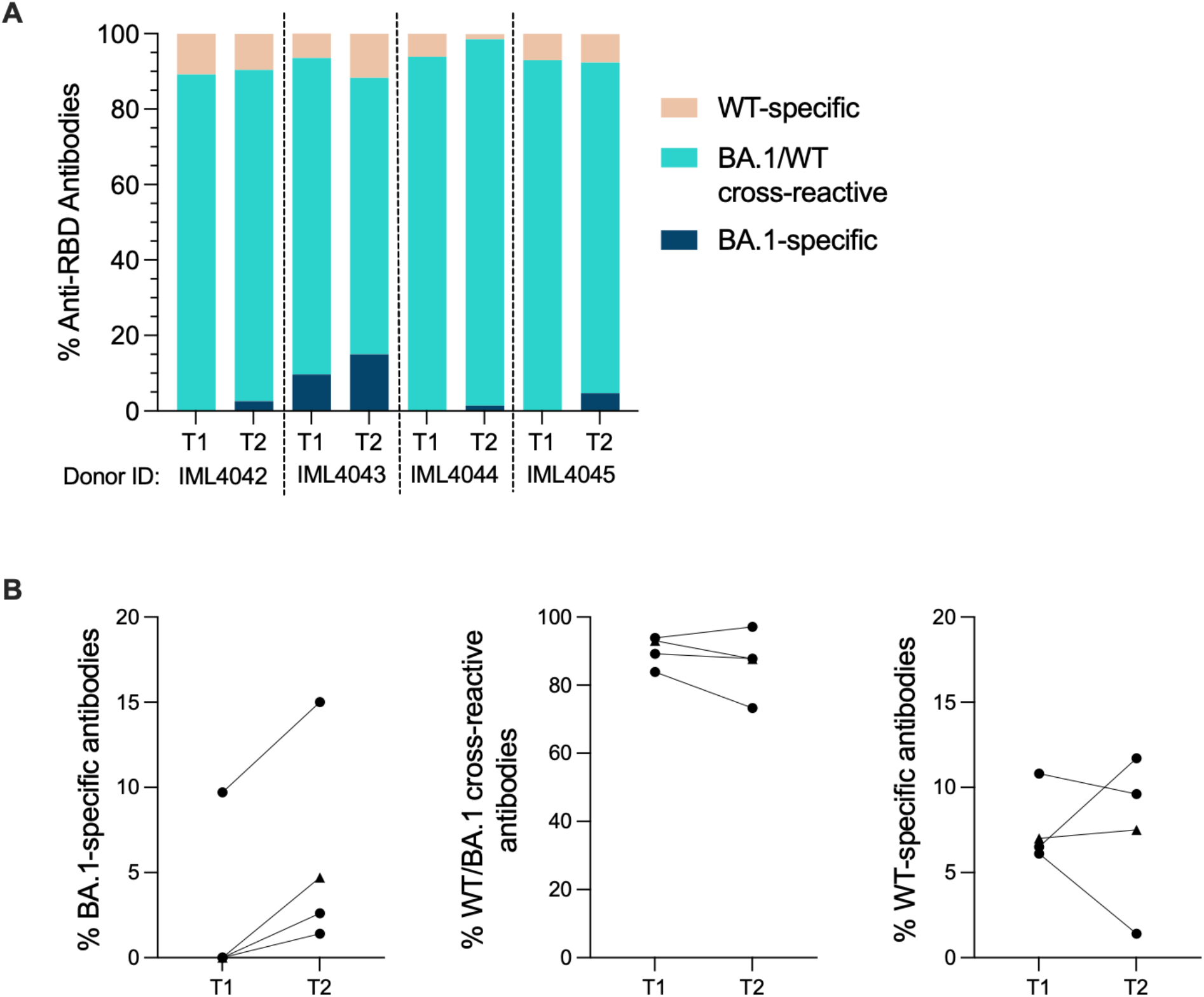
Cross-reactivity of RBD-directed monoclonal antibodies isolated at 1- and 5-6 months following BA.1 breakthrough infection. **(A-B)** Proportion of BA.1-specific, WT-specific, and WT/BA.1 cross-reactive antibodies isolated from breakthrough infection donors at 1-month (T1) and 5-6-month (T2) time points, as determined by BLI. **(B)** Summary of antibody cross-reactivity at both time points. Connected data points represent paired samples for each donor. Donors infected after two-dose mRNA vaccination (n = 4) are shown as circles and those infected after a third mRNA booster dose (n = 3) are shown as triangles. One two-dose vaccinated breakthrough donor was censored at the second time point.

**Fig. S4.**
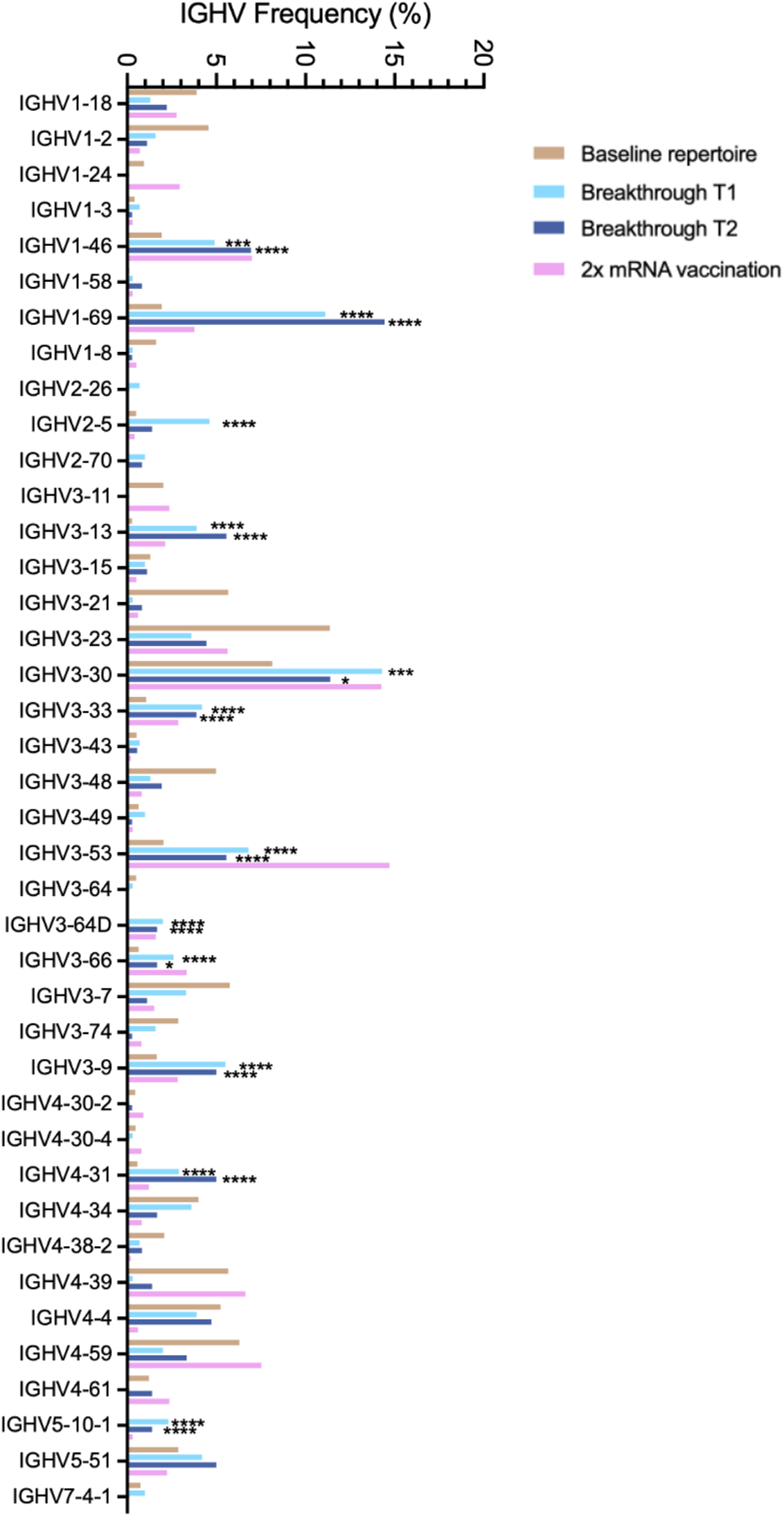
IGHV germline usage among cross-reactive antibodies. Human IGHV germline gene usage frequencies among WT/BA.1 cross-reactive antibodies at 1 month (T1) and 5-6 month (T2) time points. Germline gene distribution of RBD-directed antibodies derived from two-dose mRNA-vaccinated/uninfected donors were obtained from the CoV-AbDab database (*29*). Human baseline (unselected) repertoire frequencies were included for reference (*25*). Statistical comparisons were made by Fisher’s exact test compared to the baseline repertoire. IGHV, immunoglobulin heavy variable domain. \**P* < 0.05, \*\**P* < 0.01, \*\*\**P* < 0.001, \*\*\*\**P* < 0.0001.

**Fig. S5.**
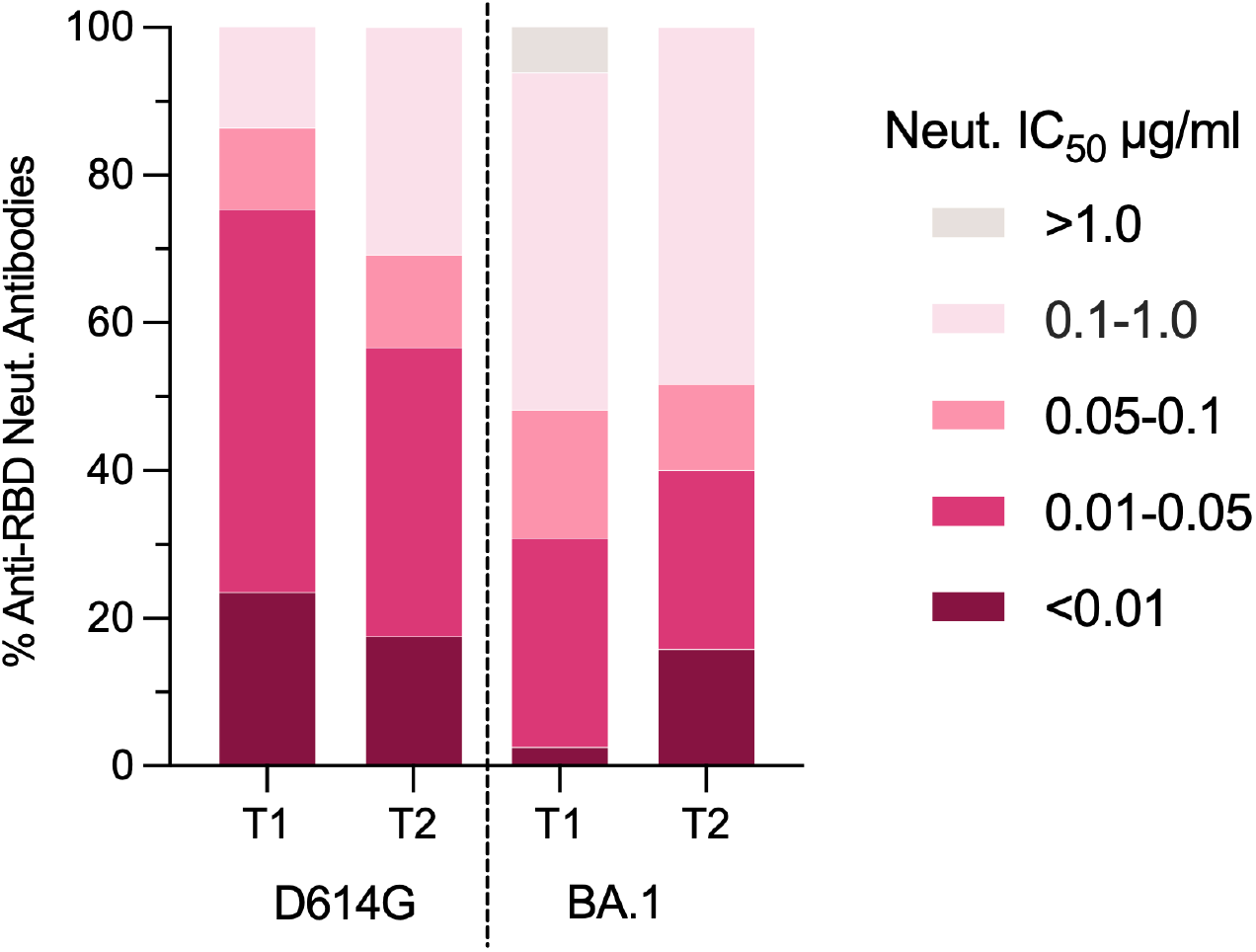
Neutralization potency of D614G/BA.1 cross-neutralizing antibodies following BA.1 breakthrough infection. Proportion of antibodies isolated at the early (T1) and late (T2) time points with the indicated neutralization IC_50_s against SARS-CoV-2 D614G and BA.1, as determined by MLV-based pseudovirus neutralization assay. Statistical comparison of BA.1 neutralizing activity by the top ten percentiles of antibodies isolated at early and late time points show significantly more potent neutralization by antibodies identified at the late time point (bootstrapping analysis of 10^th^ percentile difference using 5,000 bootstrap iterations, *P*<0.0001).

**Fig. S6.**
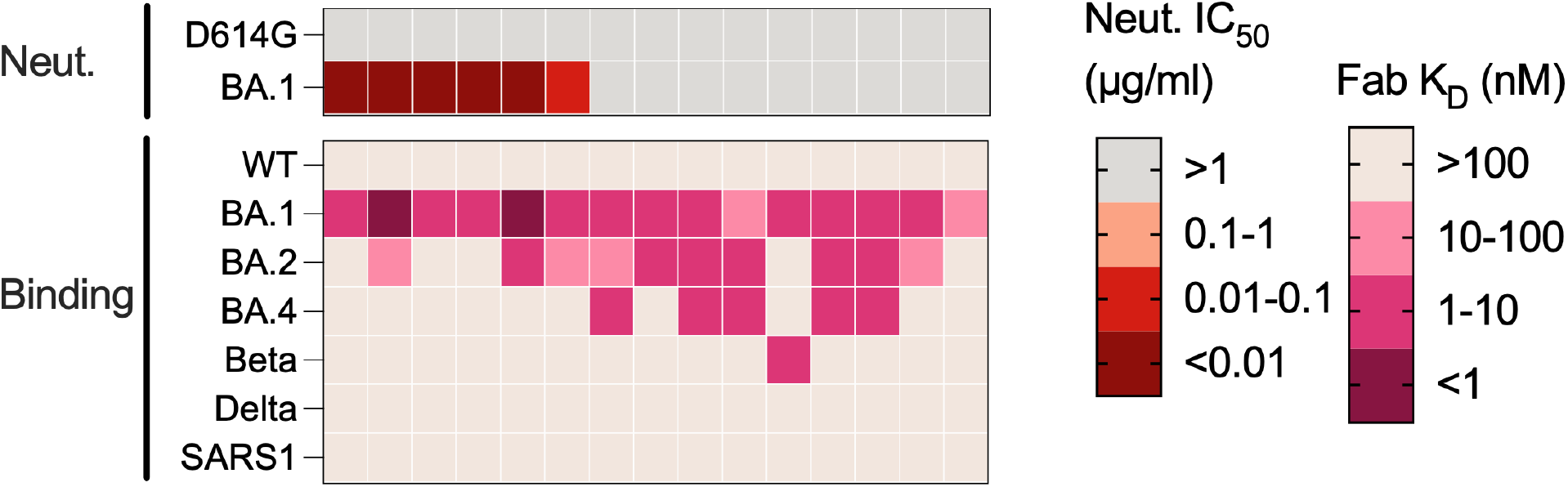
Binding and neutralization properties of BA.1-specific antibodies. Heatmap showing neutralization IC_50_s and SARS-CoV-2 variant RBD binding affinities of BA.1-specific antibodies.

**Fig. S7.**
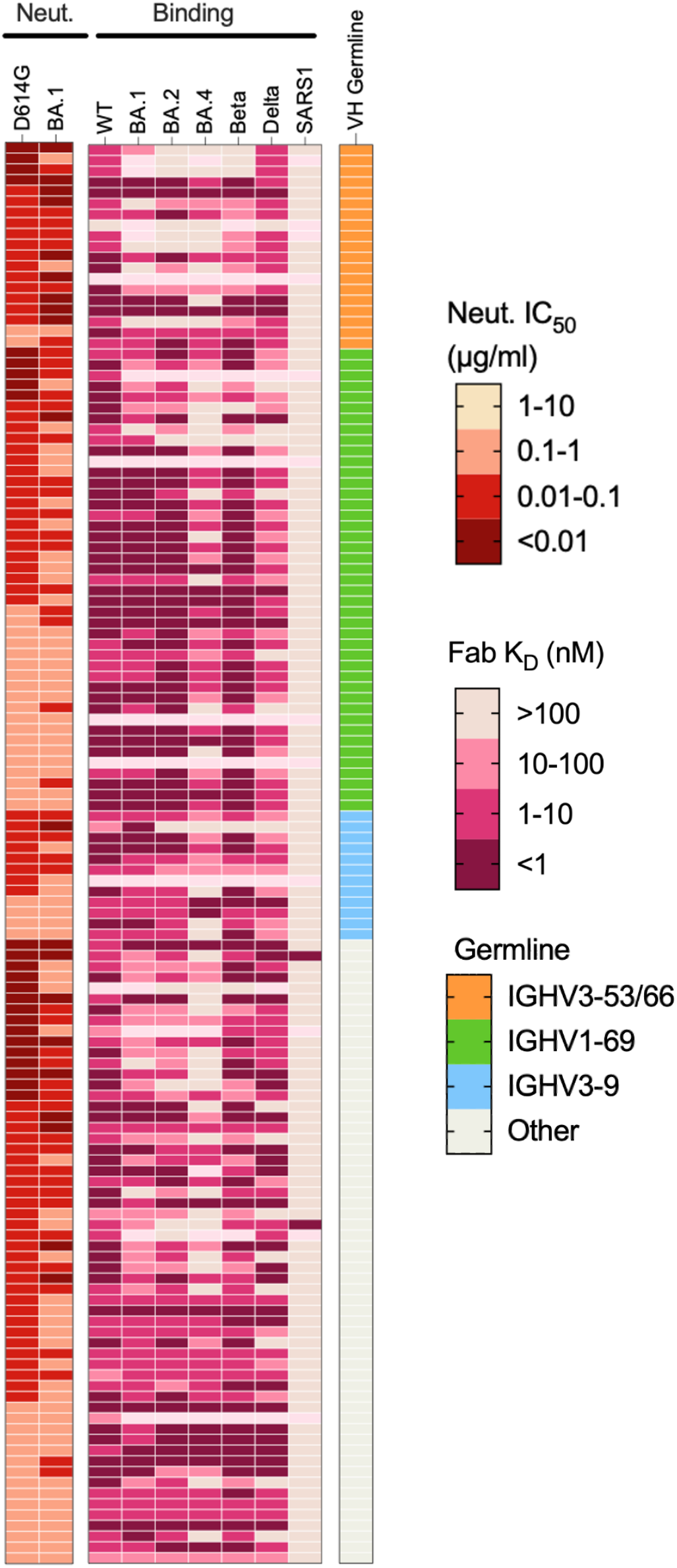
Binding breadth of D614G/BA.1 cross-neutralizing antibodies. Heatmap showing neutralization IC_50_s and SARS-CoV-2 variant RBD binding affinities of D614G/BA.1 cross-neutralizing antibodies isolated 5-6 months following BA.1 breakthrough infection. Antibodies utilizing convergent germline are indicated in the right-most column.

**Fig. S8.**
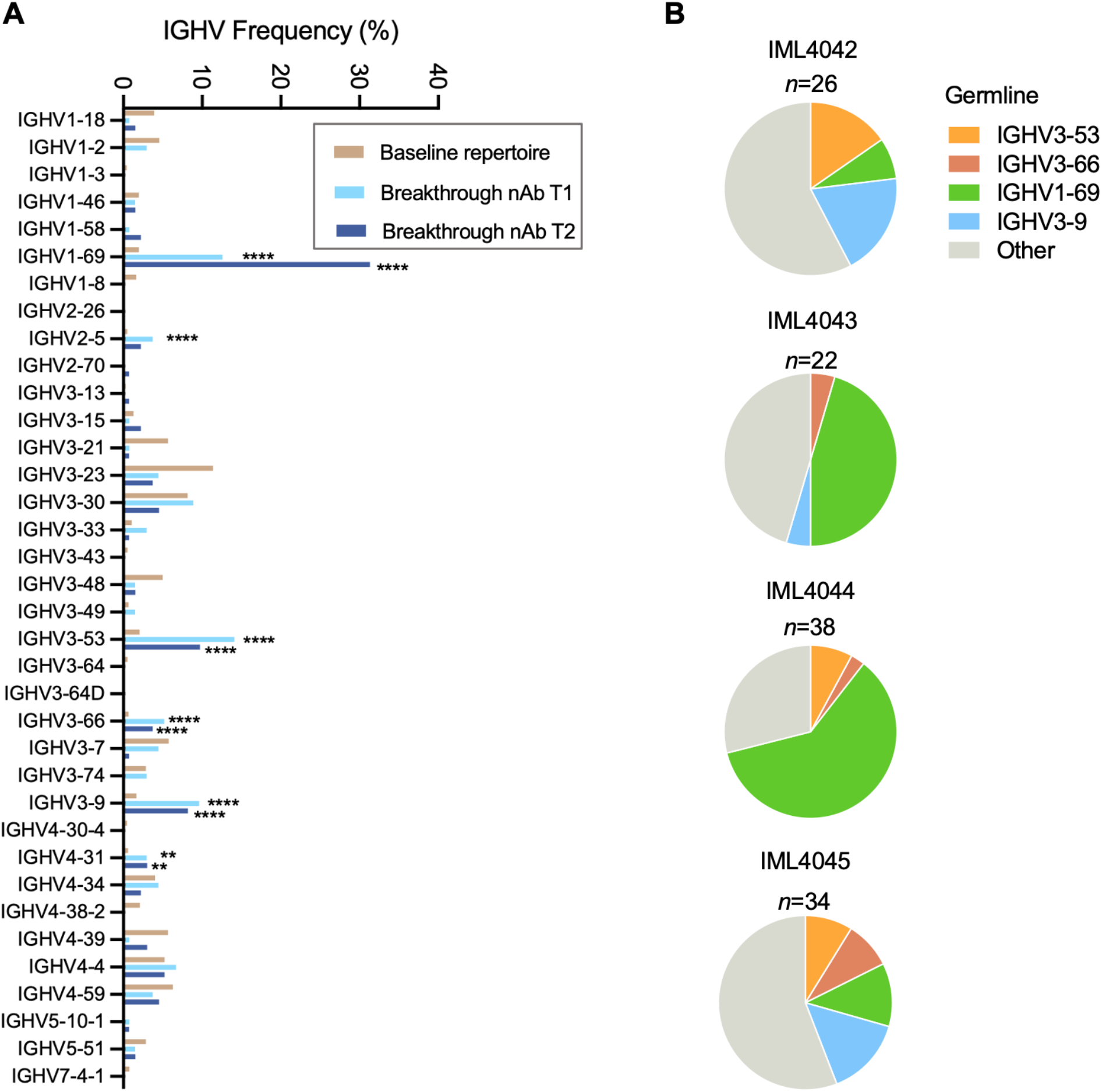
Germline gene usage of D614G/BA.1 cross-neutralizing antibodies isolated 5-6 months following BA.1 breakthrough infection. **(A)** Human IGHV germline distribution frequencies among D614G/BA.1 cross-neutralizing antibodies isolated 1-month (T1) and 5-6-months (T2) following breakthrough infection, with human baseline repertoire frequencies shown for comparison (*25*). **(B)** Pie charts showing the proportion of cross-neutralizing antibodies isolated from each donor that utilize convergent germline genes. The total number of antibodies isolated from each donor is indicated above each pie chart. IGHV, immunoglobulin human variable domain; \**P* < 0.05, \*\**P* < 0.01, \*\*\**P* < 0.001, \*\*\*\**P* < 0.0001.

**Fig. S9.**
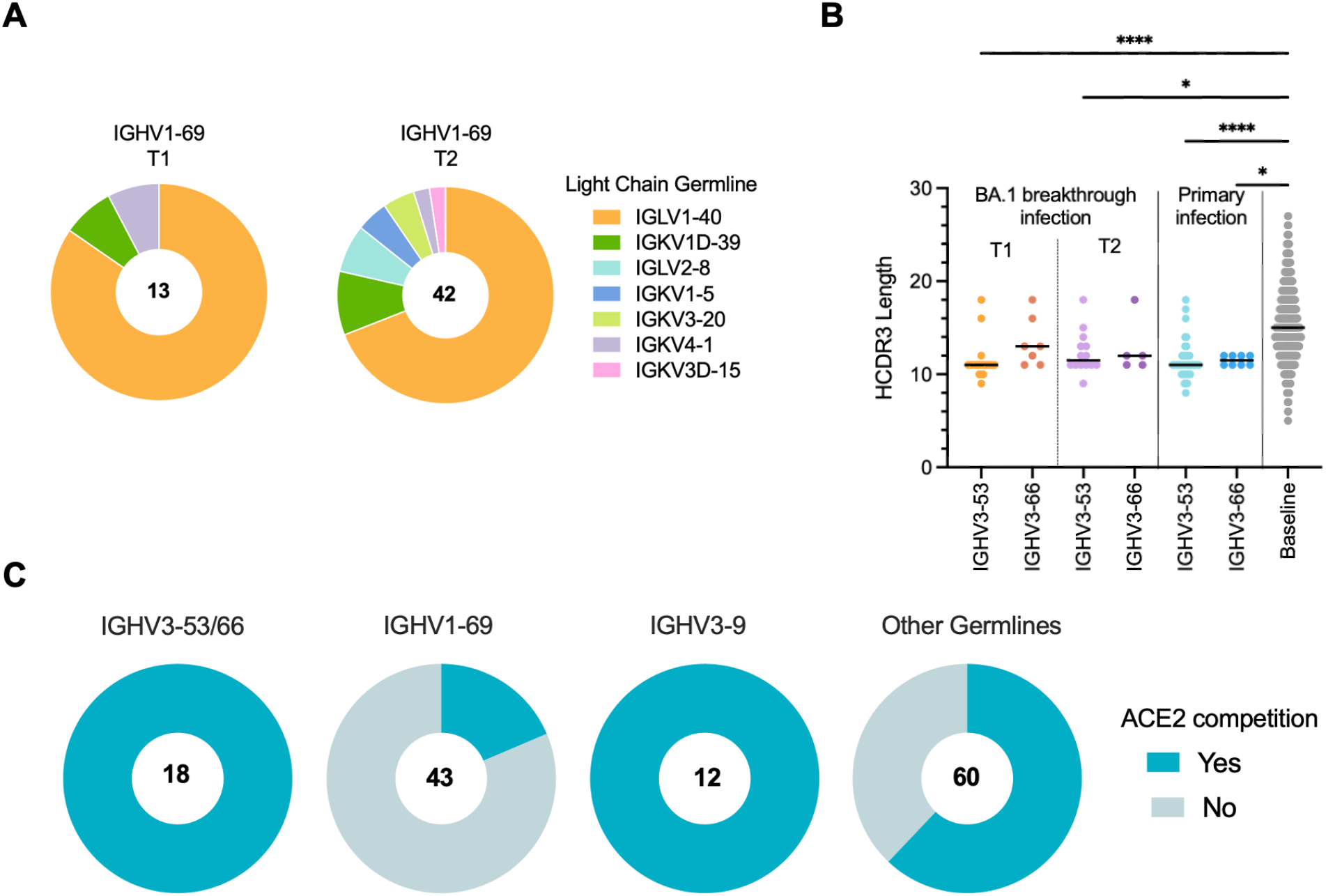
Sequence and binding features of antibodies utilizing convergent germline genes. **(A)** Pie charts showing light chain germline usage among *IGHV1-69* antibodies isolated at 1-month (T1) and 5-6 month (T2) time points. The number of antibodies analyzed from each time point is indicated in the center of each pie. **(B)** HCDR3 amino acid length distribution of *IGHV3-53* and *IGHV3-66* cross-neutralizing antibodies isolated 1-month (T1) and 5-6 months (T2) following BA.1 breakthrough infection. HCDR3 lengths of *IGHV3-53/3-66*-utilizing antibodies isolated following primary D614G infection and the baseline human antibody repertoire were included for comparison (*25*, *30*). **(C)** Proportion of cross-neutralizing antibodies utilizing the indicated germline genes that compete with the ACE2 receptor for binding, as determined by a BLI competition assay. The number of antibodies analyzed is shown in the center of each pie. Statistical comparisons were determined by Kruskal-Wallis test with subsequent Dunn’s multiple comparisons. A.A., amino acids; \**P* < 0.05; \*\*\*\**P* < 0.0001.

**Fig. S10.**
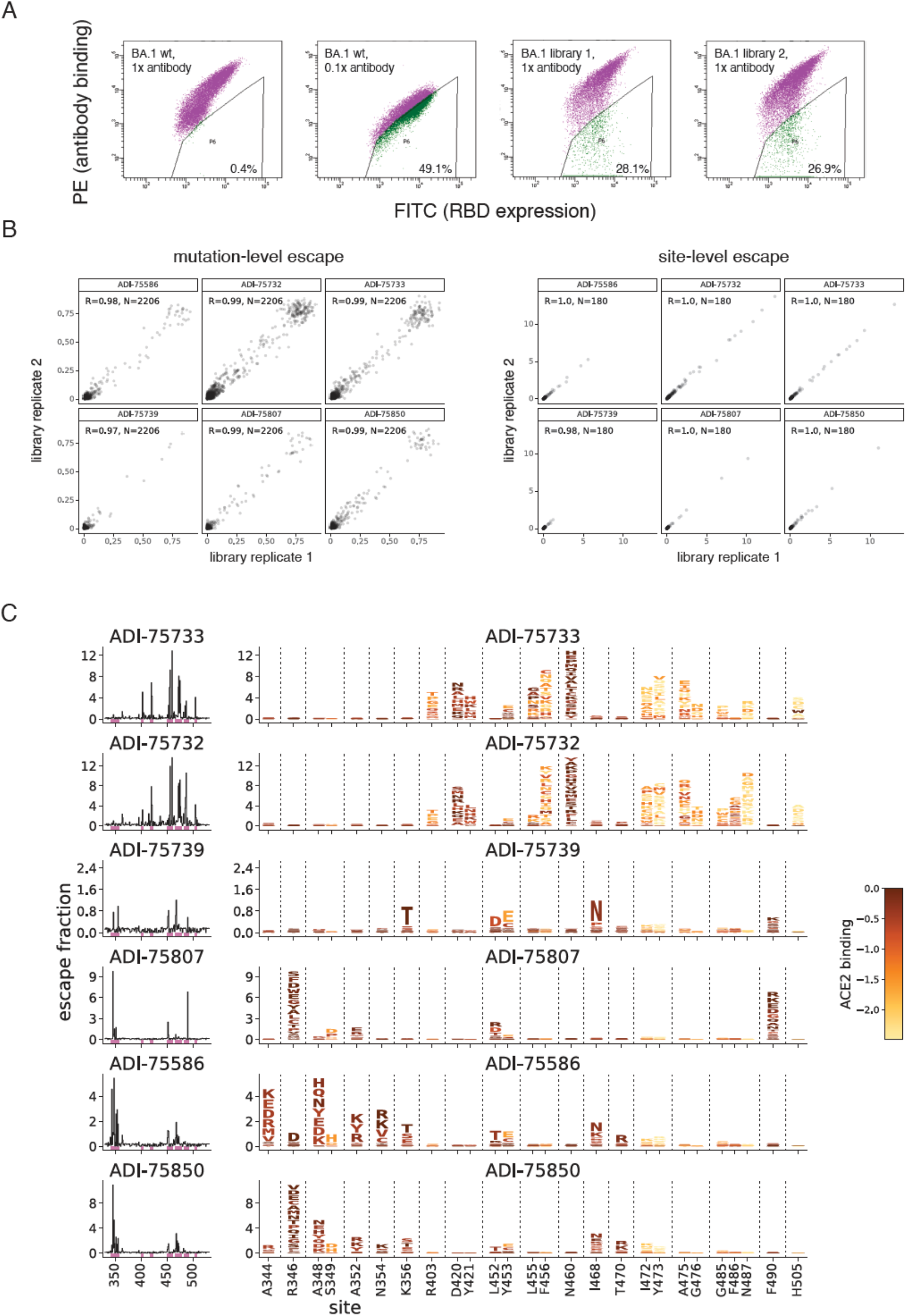
Deep mutational scanning analysis. (A) Representative FACS gates used to select antibody-escape mutations in yeast-displayed Omicron BA.1 mutant libraries. Gates were drawn to capture ~50% of wildtype Omicron BA.1-expressing yeast labeled at an antibody concentration 0.1x the selection concentration. From duplicate mutant libraries, yeast cells in the antibody-escape bin were sorted and sequenced. Post-sort mutant frequencies were compared to the pre-sort population to calculate per-mutant “escape fractions”, the fraction of cells expressing a mutation that were found in the antibody-escape sort gate. (B) Correlation in per-mutation (left) and per-site (right) escape fractions in replicate library selections for each antibody. (C) Lineplots at left show the total site-wise escape at each RBD site. This metric is mapped to structure in Fig. 3E. Sites of strong escape indicated by pink bars are shown at the mutation level in logoplots at center. Mutations are colored by their effects on ACE2 binding (scale bar at right). Note that prominent escape mutations such as K356T and I468N introduce N-linked glycosylation motifs.

---

**Table S1.** Donor Characteristics.

| Donor ID | IML4041 | IML4042 | IML4043 | IML4044 | IML4045 | IML4054 | IML4055 |
| --- | --- | --- | --- | --- | --- | --- | --- |
| Age | 45 | 19 | 23 | 23 | 24 | 38 | 23 |
| Sex | F | F | M | F | F | F | F |
| Vaccination History | 2x BNT162b2 | 2x BNT162b2 | 2x BNT162b2 | 2x BNT162b2 | 2x BNT162b2, 1x mRNA-1273 | 3x mRNA-1273 | 3x BNT162b2 |
| Date of 2nd vaccination dose | 7-May-21 | 22-Jul-21 | 23-May-21 | 10-Feb-21 | 15-May-21 | 5-May-21 | 1-May-21 |
| Date of 3rd dose (if applicable) | - | - | - | - | 20-Dec-21 | 11-Dec-21 | 9-Dec-21 |
| Date of infection | 31-Dec-21 | 4-Jan-22 | 30-Dec-21 | 2-Jan-22 | 6-Jan-22 | 19-Jan-22 | 6-Jan-22 |
| Days between infection and first (T1) sample collection | 25 | 21 | 26 | 23 | 19 | 14 | 27 |
| Days between infection and second (T2) sample collection | N/A; censored | 170 | 139 | 139 | 168 | 122 | 168 |

